# iPSC-Derived Microglia-like Cells Exhibit Protocol-Dependent Transcriptomic Features and Robust Phagocytosis of Glioma Cells

**DOI:** 10.64898/2026.07.21.739939

**Authors:** Maya N. Walker, Hui Tang, Caylee Silvers, Sasha Roth, Deanna Tiek, Bo Hu, Shi-Yuan Cheng, Xiao Song

## Abstract

Microglia are the brain-resident macrophages and key regulators of the brain tumor microenvironment. Although induced pluripotent stem cell-derived microglia (iMG) provide a valuable model for studying human microglial, systematic comparisons of differentiation protocols are limited, and their utility for modeling microglia-tumor cell interactions remains underexplored. Here, we analyzed 54 public RNA-seq datasets representing 22 iMG differentiation protocols, including embryoid body (EB)-based, two-dimensional (2D), transcription factor-induced, and coculture-based approaches. Most iMG closely resembled primary human microglia, although substantial protocol-dependent differences were observed. iMG generated using EB-based protocols showed higher *TMEM119* expression, whereas those generated using 2D-based protocols showed higher *P2RY12* expression. A widely adopted EB-based protocol showed the highest phagocytosis gene signature. Using this protocol, we generated iMG that efficiently phagocytosed patient-derived glioma stem-like cells and upregulated inflammatory and immunoregulatory genes following phagocytosis. These findings provide a transcriptomic benchmark for current iMG models and support their use in investigating microglia-glioma interactions.

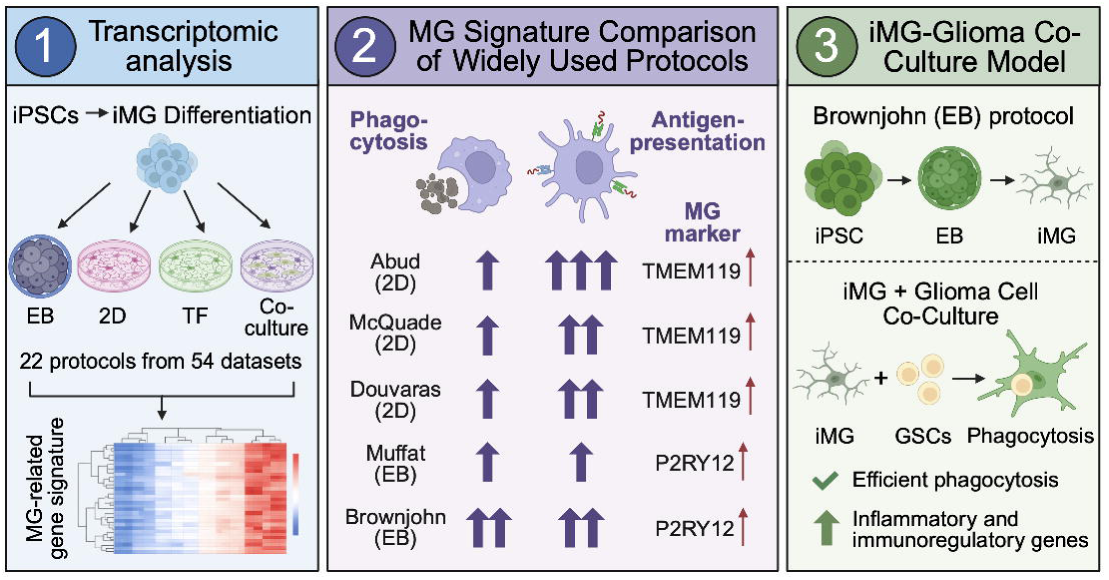

## Introduction

Microglia are the resident macrophages of the central nervous system (CNS) and play essential roles in tissue homeostasis, immune surveillance, synaptic remodeling, and phagocytosis (Colonna and Butovsky, 2017; Prinz et al., 2021). In response to pathological stimuli, microglia undergo dynamic phenotypic and functional changes, contributing to the development and progression of a wide range of neurological disorders, including neurodegenerative diseases, neuroinflammation, and CNS tumors (Khan et al., 2023; Song and Colonna, 2018). However, studies of human microglia have been limited by the restricted availability of primary brain tissue and the difficulty of maintaining primary microglia ex vivo.

Recent advances in induced pluripotent stem cell (iPSC) technology have enabled the generation of microglia-like cells that recapitulate many features of primary human microglia (Tiwari et al., 2025; Walsh and Lukens, 2025; Woolf et al., 2025). Compared with immortalized microglial cell lines such as HMC3, iPSC-derived microglia more closely resemble primary human microglia in their gene expression, inflammatory responses, and phagocytic function (Woolf *et al*., 2025). Multiple differentiation strategies have been developed, including embryoid body (EB)-based protocols, two-dimensional (2D) monolayer approaches, transcription factor (TF)-directed differentiation, and coculture systems with neural lineage cells (Tiwari *et al*., 2025). Among these, EB-based and 2D protocols are the most widely used. EB-based methods recapitulate early hematopoietic development through three-dimensional embryoid body formation (Brownjohn et al., 2018; Muffat et al., 2016; Trudler et al., 2021), whereas 2D approaches induce microglial differentiation directly in monolayer culture (Abud et al., 2017; Douvaras et al., 2017; McQuade et al., 2018). TF-directed differentiation accelerates lineage specification through the forced expression of key TFs (Chen et al., 2021; Sonn et al., 2022), while coculture systems promote microglial maturation by providing neural environmental cues (Andreone et al., 2020; Pandya et al., 2017). Together, these platforms provide scalable and genetically tractable sources of human microglia. However, substantial differences among the resulting microglia in differentiation efficiency, maturation state, transcriptional identity, and functional properties across protocols make it difficult to determine which approach most faithfully recapitulates physiological properties of human microglia.

Gliomas, the most common and malignant primary CNS tumors in adults, are characterized by an immunosuppressive microenvironment enriched with recruited resident microglia and infiltrating macrophages, which together can account for up to 30–50% of the tumor mass (Khan *et al*., 2023; Xuan et al., 2021). Through extensive bidirectional crosstalk with tumor cells, these tumor-associated microglia and macrophages contribute to tumor progression, immune evasion, and therapeutic resistance (Chen et al., 2025; Khan *et al*., 2023). Although iPSC-derived microglia (iMG) models have been extensively exploited for studies of neurodevelopmental and neurodegenerative disorders (Chadarevian et al., 2025; Douvaras et al., 2024; Heiss et al., 2025; Luo and Sugimura, 2024; Walsh and Lukens, 2025), their use in glioma research remains relatively limited. Establishing robust and physiologically relevant iMG models would facilitate mechanistic studies of microglia-glioma interactions and provide a tractable platform for investigating tumor-associated microglial functions.

In this study, we performed a comprehensive transcriptomic analysis of 54 publicly available RNA-sequencing (RNA-seq) datasets generated using 22 distinct iMG differentiation protocols (summarized in **Table 1**). Based on these analyses, we selected an EB-based protocol (Brownjohn *et al*., 2018), which showed the highest expression of phagocytosis-related genes, to investigate iMG phagocytosis of glioma tumor cells. We demonstrated that iMG generated using this protocol efficiently phagocytose three independent patient-derived glioma stem-like cell (GSC) lines and exhibit inflammatory and immunoregulatory responses following engulfment of GSCs. Our findings provide a framework for selecting appropriate iMG models and support their application in studies of microglia-glioma interactions.

## Results

### Comparative transcriptomic analysis of published iMG models

To systematically characterize the transcriptional features of iMG models, we collected all publicly available human iMG RNA-seq datasets deposited in the NCBI Gene Expression Omnibus (GEO) before April 1, 2025. A total of 54 datasets generated using 22 distinct differentiation protocols were identified, including embryoid body (EB)-based (n = 5), two-dimensional (2D) differentiation (n = 7), transcription factor (TF)-induced (n = 5), and coculture-based approaches (n = 5). To provide reference populations, we additionally analyzed RNA-seq data from primary human microglia, primary human monocytes, undifferentiated iPSCs, a human monocytic cell line THP-1, and a human microglial cell line HMC3. Details of all datasets included in this study are summarized in **Table 1** and **Table 2**. The computational analysis pipeline is summarized in **Fig. S1**.

We next examined the expression of genes associated with key macrophage and microglial functions, including phagocytosis, antigen presentation, and co-stimulatory signaling, together with a curated microglia-specific gene set derived from an integrative analysis of primary human microglia and human brain single-cell RNA-sequencing dataset (see Methods). Hierarchical clustering of iMG samples based on the expression of these genes revealed distinct transcriptomic patterns across the datasets (**Fig. 1**). Overall, iMG generated by most differentiation protocols exhibited transcriptional profiles similar to those of primary human microglia. Canonical microglial markers, including *C1QC*, *C1QB*, *C1QA*, *GPR34*, *P2RY12*, and *TMEM119*, were robustly expressed in iMG generated using most differentiation protocols, whereas these genes were expressed at substantially lower levels in monocytes, THP-1 cells, HMC3 cells, and undifferentiated iPSCs.

**Figure 1.**
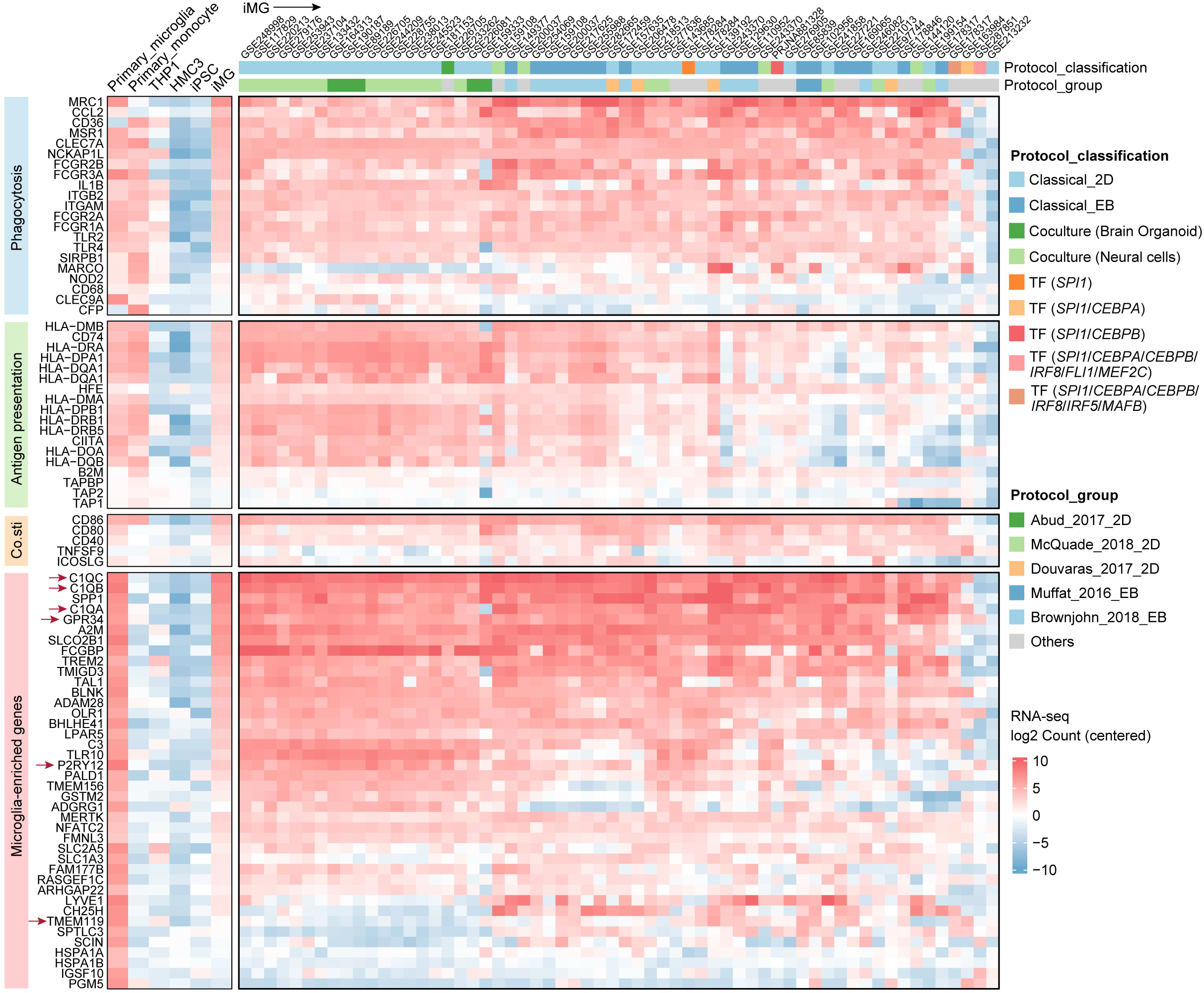
Comparative transcriptomic analysis of primary microglia, iPSC-derived microglia, and other myeloid cell models. Heatmap showing the expression of representative genes associated with phagocytosis, antigen presentation, co-stimulation, and microglial identity across human primary microglia, human primary monocytes, THP-1 cells, HMC3 cells, iPSCs, and iPSC-derived microglia (iMG). Gene expression values were log2-transformed, mean-centered, and visualized as a heatmap. iMG samples are annotated by protocol type and protocol group, with protocol groups named after the original publications. Abbreviations: iPSC, induced pluripotent stem cell; iMG, iPSC-derived microglia; 2D, two-dimension; EB, embryoid body; TF, transcription factor. Arrows highlight canonical microglial marker genes.

Despite sharing a common microglial identity, considerable heterogeneity was observed among differentiation protocols. Genes involved in phagocytosis, antigen presentation, and co-stimulatory signaling were broadly expressed across iMG generated by different differentiation protocols, although their relative expression levels varied by protocol. Notably, most EB-based protocols-produced iMG exhibited relatively higher expression of phagocytosis-associated genes, whereas most 2D protocols-produced iMG tended to display higher expression of antigen-presentation genes (**Fig. 1**). Collectively, these findings indicate that currently available iMG models closely resemble primary human microglia at the transcriptomic level, while maintaining protocol-dependent differences in genes associated with microglial functions.

### Selection of five representative iMG differentiation protocols

To further investigate protocol-dependent transcriptomic differences, we focused on protocols that were represented by at least two independent datasets for detailed comparison. These included the 2D protocols developed by Abud et al. (hereafter referred to as Abud_2017_2D) (Abud *et al*., 2017), McQuade et al. (hereafter referred to as McQuade_2018_2D) (McQuade *et al*., 2018), and Douvaras et al. (hereafter referred to as Douvaras_2017_2D) (Douvaras *et al*., 2017), as well as the EB-based protocols reported by Muffat et al. (hereafter referred to as Muffat_2016_EB) (Muffat *et al*., 2016) and Brownjohn et al. (hereafter referred to as Brownjohn_2018_EB) (Brownjohn *et al*., 2018). Although these protocols share common stages of hematopoietic and myeloid differentiation, they differ substantially in the timing of differentiation, intermediate cell populations, and cytokine combinations used to promote microglial maturation, as summarized in **Fig. 2**.

**Figure 2.**
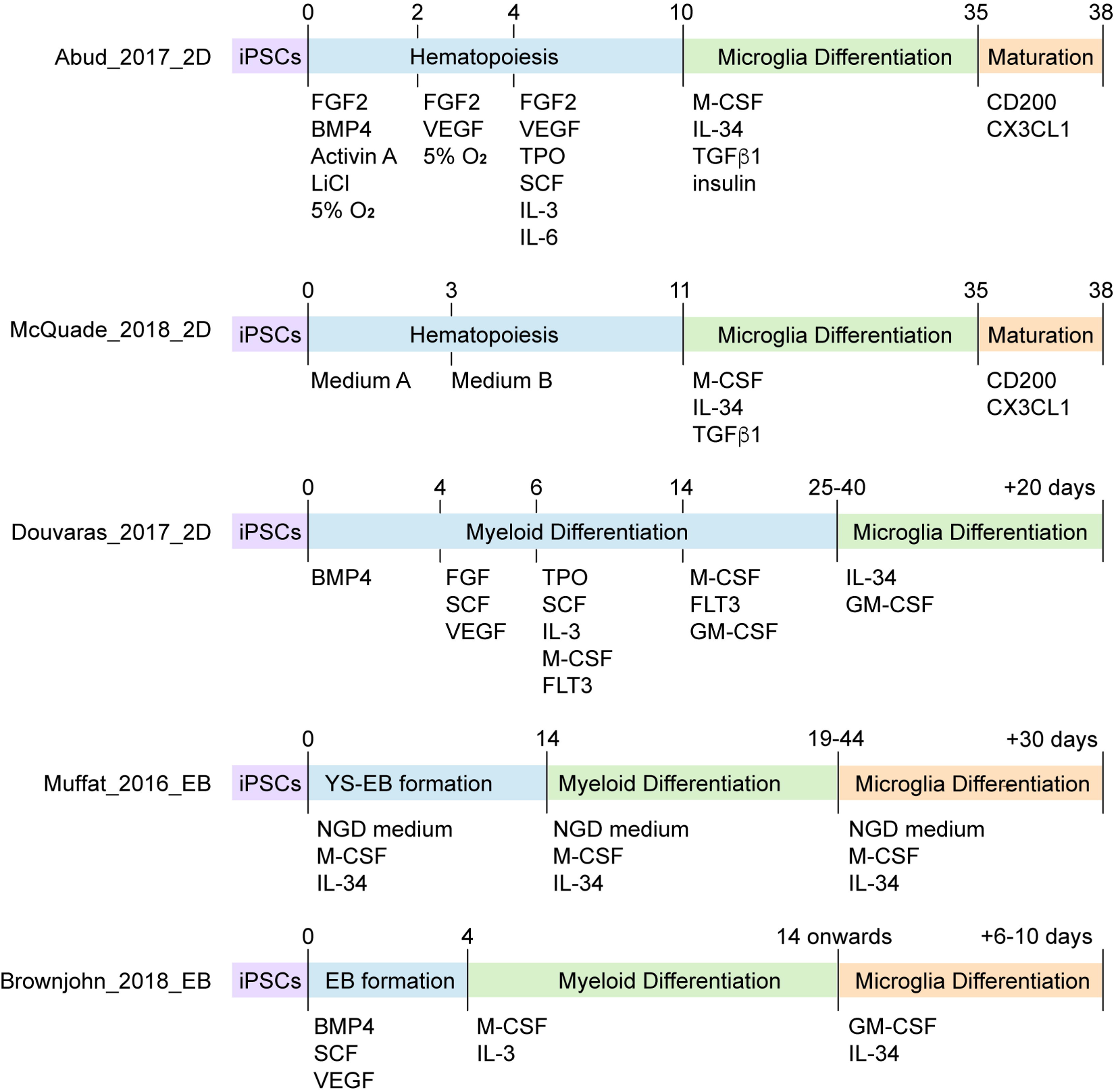
Overview of five representative iMG differentiation protocols. Schematic comparison of five representative iMG differentiation protocols representing the major protocol groups identified in the literature. Protocols include three classical 2D methods (Abud_2017_2D, McQuade_2018_2D, and Douvaras_2017_2D) and two EB-based methods (Muffat_2016_EB and Brownjohn_2018_EB). The timelines illustrate the major differentiation stages, including hematopoiesis, yolk sac embryoid body (YS-EB) formation, embryoid body (EB) formation, myeloid differentiation, microglial differentiation, and maturation, together with the corresponding cytokines, growth factors, and culture conditions used at each stage. Numbers above each timeline indicate the approximate culture day at which each differentiation stage begins. Abbreviations: iPSC, induced pluripotent stem cell; iMG, iPSC-derived microglia; 2D, two-dimensional; EB, embryoid body; YS-EB, yolk sac-like embryoid body; BMP4, bone morphogenetic protein 4; FGF2, fibroblast growth factor 2; VEGF, vascular endothelial growth factor; TPO, thrombopoietin; SCF, stem cell factor; IL, interleukin; M-CSF, macrophage colony-stimulating factor; GM-CSF, granulocyte-macrophage colony-stimulating factor; FLT3, Fms-like tyrosine kinase 3 ligand; TGFβ1, transforming growth factor-β1; CX3CL1, C-X3-C motif chemokine ligand 1.

### Protocol-dependent differences in myeloid signatures and microglial-enriched gene expression

We next compared representative iMG differentiation protocols by calculating signature scores for phagocytosis, antigen presentation, and co-stimulatory signaling in the resulting iMG (**Fig. 3**). The Brownjohn_2018_EB protocol-produced iMG exhibited the highest phagocytosis signature score among the five protocols analyzed. In contrast, all three 2D protocols-produced iMG showed relatively high antigen presentation signature scores, whereas the Muffat_2016_EB protocol-produced iMG displayed markedly lower antigen presentation activity, suggesting a reduced capacity to recapitulate the antigen-presenting properties of primary human microglia. Co-stimulatory signaling was most strongly enriched in iMG produced by Douvaras_2017_2D and Brownjohn_2018_EB protocols, indicating enhanced immune activation potential in these protocols.

**Figure 3.**
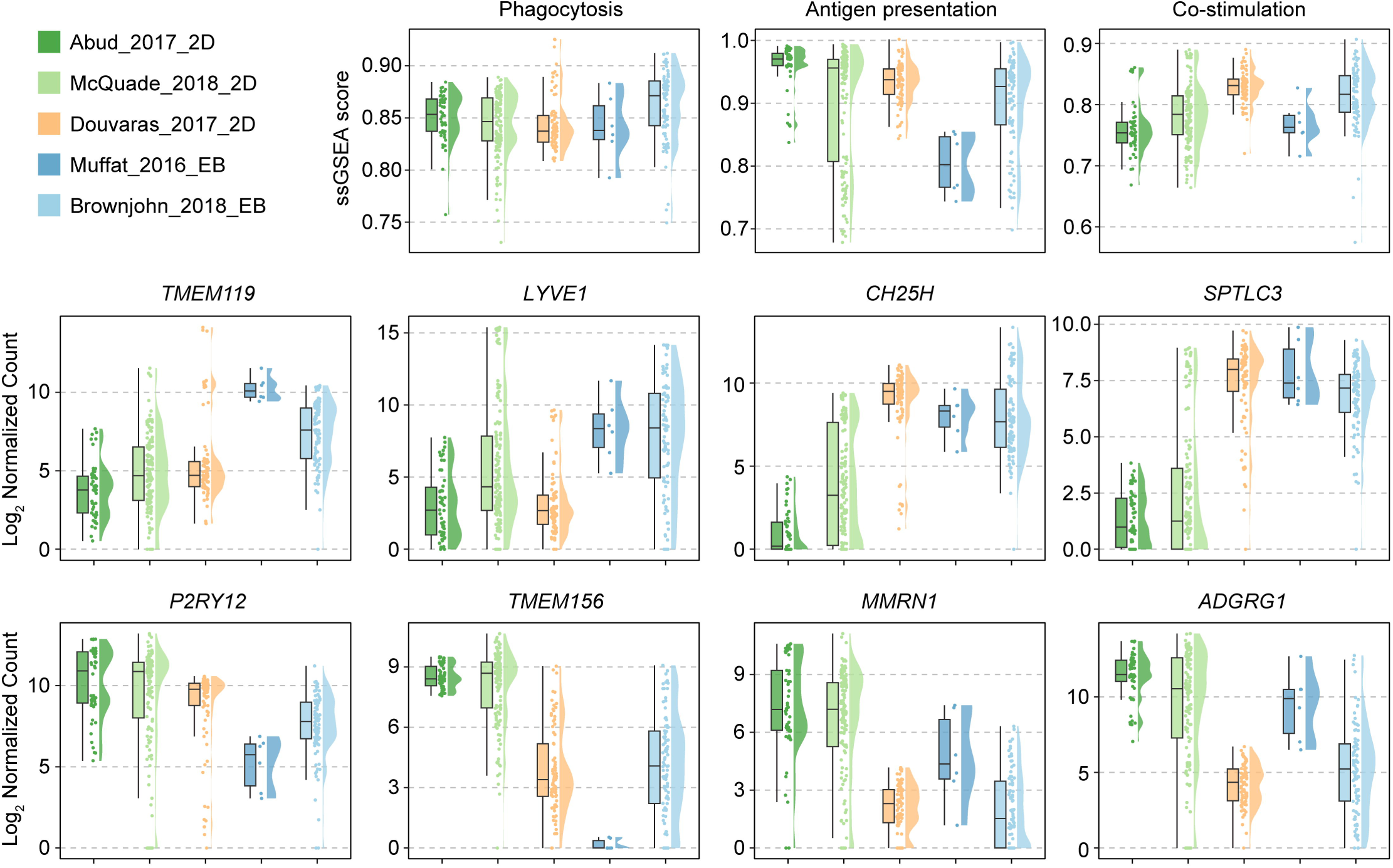
Comparison of functional gene signatures and microglial-enriched gene expression across representative iMG protocols. Single-sample gene set enrichment analysis (ssGSEA) scores for phagocytosis, antigen presentation, and co-stimulation gene signatures (top row), together with the normalized expression of representative microglial-enriched genes (bottom two rows) across five representative iMG differentiation protocols. Boxplots show the median and interquartile range, with whiskers indicating 1.5 × the interquartile range. Individual RNA-seq samples are overlaid as dots, and violin plots illustrate the distribution of ssGSEA score or gene expression (log2-transformed normalized count) within each protocol. Abbreviations: ssGSEA, single-sample gene set enrichment analysis; iMG, induced pluripotent stem cell-derived microglia; 2D, two-dimensional; EB, embryoid body.

Comparison of microglial-enriched genes involved in microglial identity and function also revealed considerable heterogeneity across differentiation protocols (**Fig. 3**). Notably, the two canonical homeostatic microglial markers *TMEM119* and *P2RY12* exhibited opposite expression patterns, with *TMEM119* preferentially expressed in EB-derived iMG, whereas *P2RY12* was consistently higher in iMG produced by the three 2D protocols. Similar protocol-dependent differences were also observed for other microglial-enriched genes, including the microglial identity markers T*MEM156*, *LYVE1*, and *MMRN1*, the immune regulator *CH25H*, the sphingolipid biosynthesis enzyme *SPTLC3*, and the adhesion G protein-coupled receptor *ADGRG1* (**Fig. 3**) Together, these findings underscore the diverse molecular phenotypes generated by different iMG differentiation protocols.

### Generation and transcriptomic characterization of iMG generated from an EB-based protocol

Based on the comparative transcriptomic analysis, we selected the Brownjohn_2018_EB protocol to model interactions between iMG and glioma tumor cells because it showed the strongest phagocytosis gene signature among the five protocols analyzed (**Fig. 4A**). Human iPSCs were differentiated into embryoid bodies and subsequently into primitive macrophage progenitors (PMPs), which were further matured into iMG in the presence of N2 supplement, IL-34, and GM-CSF. Morphological changes consistent with differentiation were observed throughout the transition from iPSCs to embryoid bodies, PMPs, and mature iMG (**Fig. 4B**).

**Figure 4.**
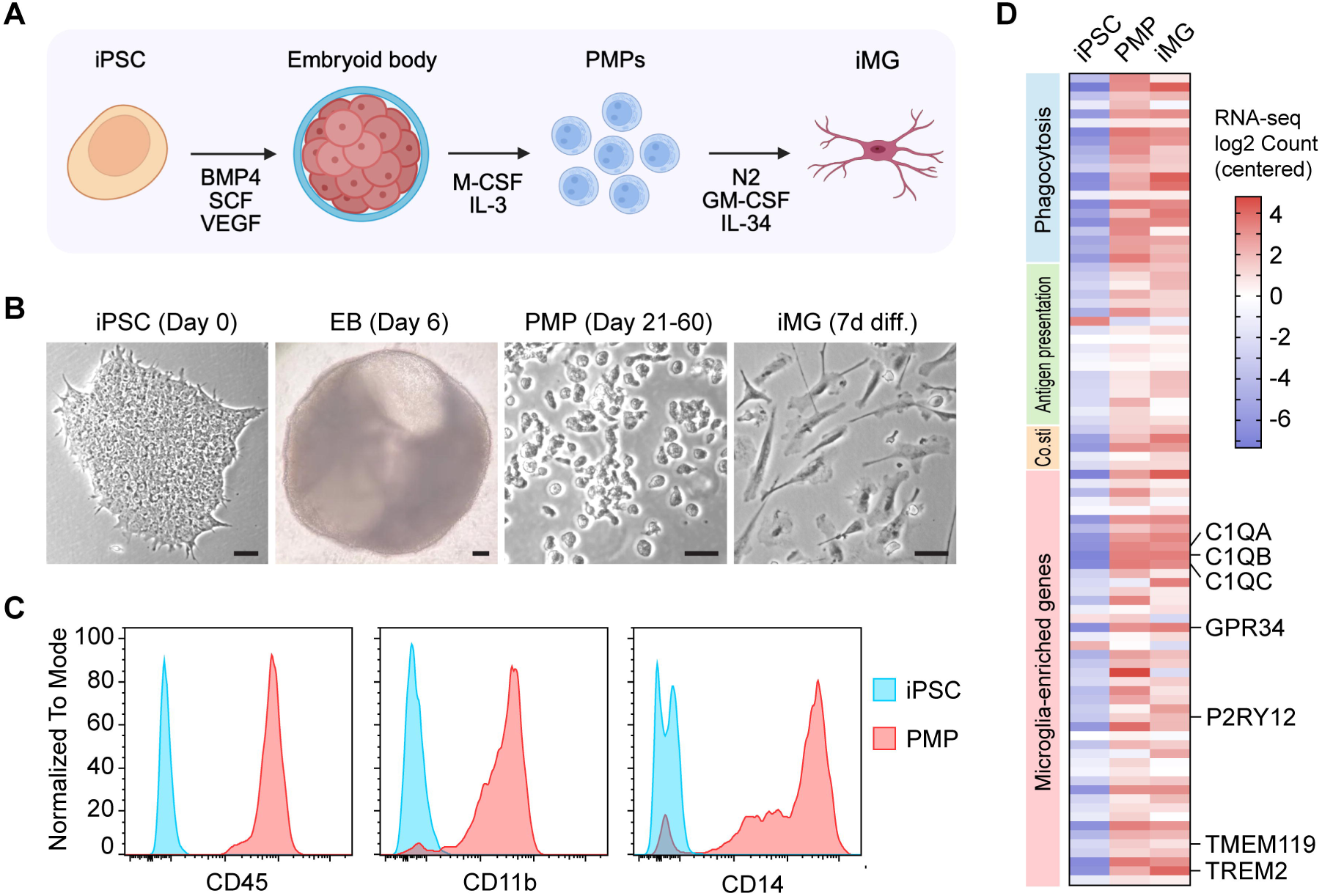
Generation and characterization of iMG using the Brownjohn EB differentiation protocol. **(A)** Schematic overview of the EB-based differentiation protocol for generating iMG. iPSCs were sequentially differentiated into embryoid bodies (EBs), primitive macrophage progenitors (PMPs), and mature iMG through stepwise cytokine induction. **(B)** Representative bright-field images of iPSCs (Day 0), EBs (Day 6), PMPs (Days 21–60), and iMG after 7 days of differentiation. Scale bars, 50 μm. **(C)** Flow cytometric analysis showing the expression of the myeloid markers CD45, CD11b, and CD14 in iPSCs and PMPs. **(D)** Heatmap showing the expression of representative genes associated with phagocytosis, antigen presentation, co-stimulation, and microglial identity in iPSCs, PMPs, and iMG. Gene expression values were log2-transformed, mean-centered, and visualized as a heatmap.

Flow cytometric analysis confirmed efficient differentiation of PMPs, which expressed the pan-leukocyte marker CD45, the myeloid marker CD11b, and the monocyte/macrophage marker CD14 (**Fig. 4C**). Transcriptomic analysis further showed that iMG exhibited increased expression of genes associated with phagocytosis, antigen presentation, co-stimulatory signaling, and microglial identity compared with undifferentiated iPSCs (**Fig. 4D** and **Table 5**). Interestingly, many microglia-enriched genes were already induced at the PMP stage and remained highly expressed following differentiation into iMG, including *P2RY12*, *TMEM119*, *GPR34*, and C1QA/B/C. Meanwhile, several genes, such as *TREM2*, exhibited a gradual increase in expression during differentiation from iPSCs to PMPs and subsequently to iMG. Together, these findings demonstrate the successful generation of iMG with molecular features characteristic of human microglia.

### EB protocol-derived iMG efficiently phagocytose glioma cells and undergo immune activation following engulfment

To determine whether EB protocol-derived iMG possess phagocytic activity toward glioma cells, we established a co-culture assay using Fluor450-labeled iMG and Fluor670-labeled patient-derived GSC line, GSC1485 (**Fig. 5A**). GSCs were exposed to 5 Gy of irradiation prior to co-culture to model therapy-induced cellular stress. Twenty-four hours after co-culture, phagocytic activity was assessed by fluorescence microscopy and flow cytometry. Fluorescence microscopy frequently detected iMG containing engulfed glioma material, suggesting active phagocytosis (**Fig. 5B**). For flow cytometric analysis, the dyes Fluor450 and Fluor670, which covalently bind to cellular proteins containing primary amines, become progressively diluted with each cell division as the fluorescent label is distributed between daughter cells. Consequently, two distinct GSC populations with high and low fluorescence intensity were observed in the GSC1485 monoculture, corresponding to non-divided and divided cells, respectively (**Fig. 5C**, second panel from the left). Phagocytic iMG were identified as Fluor450_⁺_/Fluor670_⁺_ double-positive cells in the co-culture (**Fig. 5C**, fourth and fifth panels from the left). Flow cytometric analysis demonstrated robust uptake of glioma cells by iMG under both basal and irradiated conditions, with irradiation increasing the proportion of phagocytic iMG (**Fig. 5C** and **Fig. S2A**).

**Figure 5.**
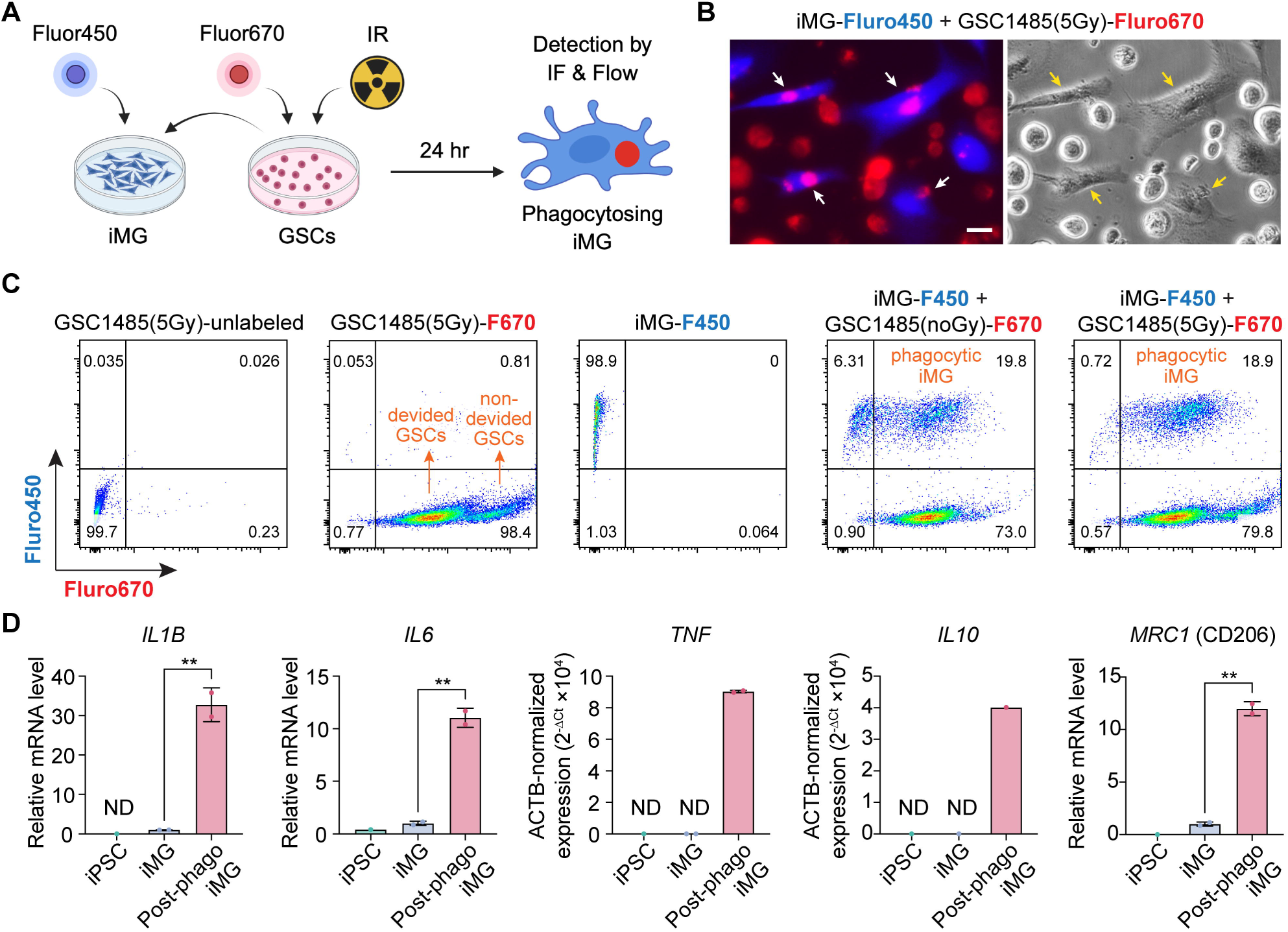
EB protocol-derived iMG efficiently phagocytose glioma cells and undergo immune activation following engulfment. **(A)** Schematic illustration of the phagocytosis assay. iMG were labeled with Fluor450 and co-cultured with Fluor670-labeled GSC1485 glioma cells following irradiation (IR, 5 Gy). After 24 hrs, phagocytosis was assessed by immunofluorescence microscopy and flow cytometry. **(B)** Representative immunofluorescence (left) and bright-field (right) images showing the uptake of Fluor670-labeled GSC1485 cells by Fluor450-labeled iMG. Arrows indicate iMG containing engulfed glioma cell. Scale bars, 20 μm. **(C)** Flow cytometry plots showing the quantification of phagocytosis. The first three panels show the control samples, including unlabeled GSC1485 cells, Fluor670-labeled GSC1485 cells, and Fluor450-labeled iMG. The last two panels show co-cultures of Fluor450-labeled iMG with Fluor670-labeled GSC1485 cells with or without 5 Gy irradiation. Double-positive (Fluor450_⁺_Fluor670_⁺_) cells represent iMG that have phagocytosed glioma cells. **(D)** Relative mRNA expression of *IL1B*, *IL6*, *TNF*, *IL10*, and *MRC1* (CD206) in iPSCs, iMG, and post-phagocytic iMG measured by quantitative PCR. Gene expression was normalized to *ACTB*. Data are presented as mean ± SD. P values were determined by two-tailed Student’s t-test. **, p < 0.01. ND, not detected.

To determine how phagocytosis influences the molecular phenotype of iMG, we isolated pre- and post-phagocytic iMG and analyzed gene expression by quantitative PCR (qPCR). Post-phagocytosis (Post-phago) iMG exhibited significantly increased expression of *IL1B* and *IL6* compared with pre-phagocytosis iMG and iPSCs (**Fig. 5D**). Increased expression of *TNF*, *IL10*, and *MRC1* (CD206) was also observed following phagocytosis, indicating activation of both inflammatory and immune-regulatory programs. These findings demonstrate that iPSC-derived microglia efficiently engulf glioma cells and undergo transcriptional changes associated with functional activation following phagocytosis.

## Discussion

Human induced pluripotent stem cell-derived microglia (iMG) have become an increasingly important model for investigating human microglial biology and disease (Luo and Sugimura, 2024; Walsh and Lukens, 2025). However, the rapid expansion of differentiation strategies with distinct variations has made it difficult to determine which protocols can most faithfully recapitulate primary human microglia and which are best suited for specific experimental applications. In this study, we performed a systematic transcriptomic analysis by comparing 54 publicly available RNA-seq datasets representing 22 iMG differentiation protocols and combined these analyses with experimental validation using an EB-based differentiation strategy. Our results demonstrate that most currently available iMG models broadly recapitulate the transcriptional characteristics of primary human microglia while exhibiting substantial protocol-dependent differences in immune-related gene expression and function.

A major finding of this study is that most differentiation protocols are capable of successfully establishing a core microglial identity. Canonical microglial markers, including *C1QA*, *C1QB*, *C1QC*, *GPR34*, *TMEM119*, and *P2RY12*, were broadly expressed across the majority of the collected iMG datasets, supporting the overall ability of current differentiation methods to generate iMG cells with microglial characteristics. Nevertheless, comparison of representative protocols-produced iMG revealed pronounced differences in genes associated with phagocytosis, antigen presentation, co-stimulatory signaling, and microglial identity. Rather than identifying a single universally superior protocol, our results suggest that individual differentiation strategies generate iMG populations with distinct functional biases. EB-based protocols generally exhibited stronger phagocytosis-associated signatures and higher expression of *TMEM119*, whereas several 2D protocols displayed enhanced antigen-presentation signatures and higher *P2RY12* expression. These observations indicate that different differentiation methods may preferentially model different aspects of microglial biology and should therefore be selected according to the intended biological question.

The observed differences among the resulting iMG likely reflect the distinct developmental trajectories used by current differentiation strategies. EB-based protocols recapitulate primitive hematopoiesis through three-dimensional embryoid body formation before generating primitive macrophage progenitors, thereby more closely resembling the yolk sac-derived developmental origin of microglia (Brownjohn *et al*., 2018; Muffat *et al*., 2016; Trudler *et al*., 2021). In contrast, 2D monolayer differentiation bypasses aspects of early embryonic development and may favor more rapid acquisition of specific immune functions (Abud *et al*., 2017; Douvaras *et al*., 2017; McQuade *et al*., 2018). Although further mechanistic studies are warranted, our findings support the concept that developmental context contributes to the maturation state and functional specialization of iPSC-derived microglia.

Using the Brownjohn EB protocol (Brownjohn *et al*., 2018) for experimental validation, we further demonstrated efficient differentiation of iPSCs into PMPs and mature iMG. Transcriptomic profiling revealed progressive acquisition of microglial-enriched genes during differentiation, with induction of many microglial genes already evident at the PMP stage and maintained following maturation into iMG. These observations suggest that aspects of the microglial transcriptional program may begin to emerge during the progenitor stage, although further studies are needed to define the timing and mechanisms of this process.

Because microglia are major cellular components of the glioma tumor microenvironment (TME) (Khan *et al*., 2023; Xuan *et al*., 2021), we next evaluated whether iMG recapitulate one of their hallmark functions—tumor cell phagocytosis. The generated iMG efficiently engulfed three independent patient-derived GSC lines, demonstrating that their phagocytic activity is reproducible across glioma models. Notably, irradiation of GSCs further enhanced phagocytosis, consistent with the possibility that therapy-induced cellular stress increases tumor cell recognition or susceptibility to engulfment by microglia (Gholamin et al., 2020). Following phagocytosis, iMG upregulated *IL1B*, *IL6*, *TNF*, *IL10*, and *MRC1* (CD206), indicating activation of both inflammatory and immunoregulatory programs. These findings demonstrate that iPSC-derived microglia not only possess robust phagocytic capacity but also mount dynamic transcriptional responses following tumor cell engulfment, supporting their utility for studying functional microglia-glioma interactions.

This study has several limitations. First, the comparative transcriptomic analyses integrated datasets generated by multiple laboratories using different sequencing platforms, differentiation protocols, and experimental conditions, all of which may contribute to inter-study variability. Although uniform bioinformatic processing minimized technical differences, some variability is likely attributable to batch effects rather than biological differences alone. Second, functional validation was performed using a single EB-based differentiation protocol. Future studies directly comparing multiple differentiation strategies under standardized culture conditions is warranted to determine whether the protocol-dependent transcriptional differences identified here translate into distinct functional phenotypes. Finally, while our in vitro assays demonstrate robust phagocytic activity and activation following tumor cell engulfment, they cannot fully recapitulate the complex cellular interaction and environmental cues present within the glioma TME in vivo.

In summary, this study provides a systematic transcriptomic comparison of currently available iMG differentiation protocols and demonstrates that protocol selection influences both microglial identity and immune-related functional programs. Our experimental validation further establishes that EB-derived iMG undergo progressive acquisition of microglial characteristics, efficiently phagocytose patient-derived glioma cells, and exhibit dynamic transcriptional activation following tumor cell engulfment. Together, these findings provide a practical framework for selecting appropriate iMG models and support the use of iPSC-derived microglia as a tractable platform for investigating microglia-glioma interactions in the TME and evaluating therapeutic strategies targeting the glioma immune microenvironment.

## Methods

### Collection of public iMG transcriptomic datasets

Publicly available human RNA-sequencing datasets were downloaded from the NCBI Gene Expression Omnibus (GEO) database. Datasets were identified using the search terms “iPSC” and “microglia” and filtered to include human studies generated by high-throughput RNA sequencing. All datasets deposited before April 1, 2025, were considered. Samples corresponding to iMG were manually curated based on sample annotations and the associated publications. A total of 54 RNA-seq datasets representing 22 distinct iMG differentiation protocols were included in the analysis. Detailed information for each iMG dataset, including GEO accession number, sample ID, original publication, differentiation method, protocol name, and protocol classification, is provided in **Table 1**. Because multiple independent studies adopted the same differentiation protocol, datasets were grouped according to the original protocol for protocol-level analyses. The 22 differentiation protocols were further classified into four major categories: embryoid body (EB)-based, two-dimensional (2D), transcription factor (TF)-directed, and coculture-based approaches involving neural lineage cells or brain organoids. RNA-seq datasets from primary human microglia, primary human monocytes, undifferentiated iPSCs, the monocytic cell line THP-1, and the microglial cell line HMC3 were additionally included as reference cell populations, and details of these datasets are provided in **Table 2**.

### RNA-sequencing data processing

Raw sequencing data were downloaded from the Sequence Read Archive (SRA) using SRA Toolkit. Sequencing quality was assessed using FastQC (v0.11.5). Reads were aligned to the human reference genome GRCh38 using STAR (v2.7.9a) (Dobin et al., 2013) in two-pass mode with GENCODE v44 gene annotations. Gene-level read counts were generated using HTSeq-count (v2.0.2) (Putri et al., 2022). Gene count matrices from all datasets were merged and normalized using DESeq2 (v1.52.0) (Love et al., 2014) for downstream analyses. All scripts used for RNA-seq data processing and downstream analyses have been deposited in Zenodo (10.5281/zenodo.21431874).

### Identification of microglia-enriched genes

To establish a reference microglia-enriched gene set, differential gene expression analyses were performed between primary human microglia and primary human monocytes, as well as between primary human microglia and undifferentiated iPSCs. Raw count matrices were analyzed using DESeq2 (v1.52.0). Genes were considered microglia-enriched if they satisfied the following criteria in both comparisons: (i) log2 fold change > 4; (ii) Benjamini–Hochberg adjusted P < 0.001; (iii) mean normalized count > 1,000 in primary human microglia; and (iv) protein-coding gene annotation.

To further increase the specificity of the microglia-enriched gene set, marker genes were identified from the Allen Brain Cell Atlas human primary motor cortex (M1) single- cell RNA-sequencing dataset (10x Genomics platform), downloaded from the Allen Brain Cell Atlas data portal (https://portal.brain-map.org/atlases-and-data/rnaseq/human-m1-10x). Raw gene expression matrices and the metadata were imported into Seurat (v5.5.1) (Hao et al., 2024), and cells were normalized using the LogNormalize method with a scaling factor of 10,000. Microglial marker genes were identified by comparing the “Micro L1-6 TYROBP CD74” cluster (microglia cluster) with all other brain cell populations using the FindMarkers function with the Wilcoxon rank-sum test. Genes with an average log2 fold change > 4 were considered microglia-enriched. The resulting marker genes were intersected with the bulk RNA-seq-derived microglia-enriched gene set to generate a high-confidence reference microglial gene signature for subsequent analyses.

### Functional gene signatures

Gene sets representing phagocytosis, antigen presentation, and co-stimulatory signaling were manually compiled based on genes with established functions in these pathways (**Table 3**). Together with the microglia-enriched gene set, these signatures were used to evaluate functional characteristics across iMG differentiation protocols. Gene set activity was quantified using single-sample Gene Set Enrichment Analysis (ssGSEA) implemented in the GSVA package (v2.6.2) (https://github.com/guokai8/scGSVA). Briefly, log2-transformed normalized expression values were used as input, and ssGSEA enrichment scores were calculated for each sample and compared across iMG differentiation protocols.

### Comparative transcriptomic analysis

Normalized gene expression values were log2-transformed for downstream analyses. For heatmap visualization, expression values were mean-centered across all samples. Gene expression values were averaged across biological replicates within each cell type or differentiation protocol. Heatmaps were generated using the ComplexHeatmap package (v2.28.0) (Gu et al., 2016) in R, and hierarchical clustering was performed using Manhattan distance and Ward’s linkage method. Expression patterns of genes associated with phagocytosis, antigen presentation, co-stimulatory signaling, and microglial identity were examined across all iMG differentiation protocols. Five representative differentiation protocols represented by at least two independent datasets were selected for detailed comparative analyses. Expression of selected genes or ssGSEA signature scores was visualized using violin plots and boxplots generated with ggplot2 (v4.0.3).

### Cell Culture

Patient-derived glioma stem-like cell (GSC) lines GSC1485, GSC11, GSC528 were previously established (Bhat et al., 2013; Song et al., 2019) and maintained as neurospheres in GSC culture medium, which includes DMEM/F12 medium (Thermo Fisher Scientific, 11320-033), 2% B27 supplement (Thermo Fisher Scientific, 17504-044), 1 × antibiotic-antimycotic (Thermo Fisher Scientific, 15240062), 5 mg/mL heparin (Sigma-Aldrich, 9041-08-1), 20 ng/mL EGF (Peprotech, 100-15R), and 20 ng/mL bFGF (Peprotech, 100-18B).

Human iPSCs (iPSC12 line) were kindly provided by Dr. Frank Funari at University of California, San Diego (Miki et al., 2022). iPS cells were cultured on plates coated with Matrigel hESC-Qualified Matrix (Corning) in mTeSR Plus medium (Stemcell Technologies). iPSC-derived NPCs were cultured on matrigel-coated plates in N2B27 medium (50% DMEM/F12 and 50% Neurobasal-A medium supplemented with GlutaMAX, 1 × N-2 supplement, 1 × B-27 supplement, 1 × antibiotic-antimycotic, 150 µM ascorbic acid) plus 3 μM CHIR99021 (DNSK International) and 1 μM SAG (DNSK International).

### Generation of iMG

Human iPSCs were differentiated into iMG using an EB-based protocol adapted from Brownjohn et al (Brownjohn *et al*., 2018; Li et al., 2024). Briefly, single-cell suspensions of iPSCs were aggregated in low-attachment plates to generate embryoid bodies in mTeSR Plus medium supplemented with BMP4 (50 ng/mL), VEGF (50 ng/mL), and SCF (20 ng/mL). After six days, embryoid bodies were transferred to primitive macrophage progenitors (PMP) medium consisting of X-VIVO 15 supplemented with non-essential amino acids (NEAA), GlutaMAX, β-mercaptoethanol, M-CSF (100 ng/mL), and IL-3 (25 ng/mL) to generate PMPs. PMPs were continuously released into the culture medium and harvested beginning approximately 2-4 weeks after EB formation. For microglial differentiation, PMPs were plated at 2 × 10^5^ cells/mL and cultured for seven days in DMEM/F12 supplemented with N2, IL-34 (100 ng/mL), GM-CSF (10 ng/mL), NEAA, and β-mercaptoethanol, with medium changes every three days. Mature iMG were subsequently used for flow cytometry, transcriptomic characterization, and functional assays.

### Flow cytometry

PMPs were harvested and stained with Human TruStain FcX™ Fc receptor blocking solution (BioLegend, Cat. No. 422302), followed by fluorophore-conjugated antibodies against human CD45-FITC (BioLegend, Cat. No. 304006), CD11b-APC (BioLegend, Cat. No. 340007), and CD14-PE (BioLegend, Cat. No. 325606). Antibodies were used according to the manufacturers’ recommended staining conditions. Data were acquired on a BD FACSymphony™ A5 Cell Analyzer (BD Biosciences) and analyzed using FlowJo (v10; BD Biosciences).

### RNA-seq analysis of iPSCs, PMPs, and iMG

Total RNA was extracted from iPSCs, primitive macrophage progenitors (PMPs), and iMG using the RNeasy Mini Kit (Qiagen) according to the manufacturer’s instructions. Libraries for Illumina sequencing were prepared using NEBNext Poly(A) mRNA Magnetic Isolation Module, Ultra II RNA Library Prep Kit, and Multiplex Oligos (NEB, Cat #E7490, E7775, E6609) according to the manufacturer’s instruction. Sequencing was performed on the Illumina NovaSeq platform using 150 bp paired-end run with 70-100 million paired reads per sample at the Northwestern University NUSeq core facility. Raw sequencing reads were processed using the same bioinformatic pipeline described above in the “RNA-sequencing data processing” section.

### Glioma cell phagocytosis assay

Patient-derived GSC1485 cells were cultured either under basal conditions or exposed to 5 Gy ionizing radiation. Irradiated cells were maintained for an additional three days before the phagocytosis assay. GSC1485 cells were labeled with eBioscience™ Cell Proliferation Dye eFluor™ 670 (Thermo Fisher Scientific, Cat. No. 65-0842-90), and iMG were labeled with eBioscience™ Cell Proliferation Dye eFluor™ 450 (Thermo Fisher Scientific, Cat. No. 65-0840-90), according to the manufacturer’s instructions. Labeled glioma cells and iMG were co-cultured at a 4:1 ratio for 24 hrs. Phagocytosis was assessed by fluorescence microscopy and flow cytometry. Phagocytic iMG were identified as eFluor™ 450_⁺_/eFluor™ 670_⁺_ double-positive cells. The phagocytic capacity of iMG was further validated using two additional patient-derived GSC lines, GSC528 and GSC11. The GSC11 line was stably transduced with a lentiviral reporter vector co-expressing firefly luciferase and the red fluorescent protein dsRed.

### Quantitative PCR

Post-phagocytosis iMG populations, defined as eFluor™ 450_⁺_/eFluor™ 670_⁺_ double-positive cells, were isolated by fluorescence-activated cell sorting following co-culture with GSCs. Total RNAs from post-phagocytosis iMG, pre-phagocytosis iMG, and iPSCs were isolated using a Qiagen RNeasy Mini Kit and were reverse transcribed using the iScript cDNA Synthesis Kit (Bio-Rad) according to the manufacturer’s instructions. Quantitative RT-PCR was performed using the EvaGreen 2 x qPCR MasterMix (Bullseye) on an Applied Biosystems StepOne Plus Real-Time Thermal Cycling Block. Relative gene expression was determined by normalizing the expression of each target gene to ACTB. Results were analyzed using the 2^-(ΔΔCt)^ method. Primers are listed in **Table 4**.

### Statistical analysis

Statistical analyses were performed in R (v4.6.1) or GraphPad Prism (v10). Differential gene expression analyses were conducted using DESeq2 with Benjamini–Hochberg correction for multiple testing, and log2 fold changes were estimated using apeglm shrinkage. For qPCR experiments, differences between post-phagocytosis and pre-phagocytosis iMG populations were evaluated using unpaired two-tailed Student’s t-tests. Data are presented as mean ± SD. Unless otherwise stated, P < 0.05 was considered statistically significant.

## Data availability

The RNA-seq datasets generated in this study are available from the corresponding author upon reasonable request.

## Code availability

All scripts used for RNA-seq processing, data integration, differential expression analysis, and downstream analyses have been deposited in Zenodo (10.5281/zenodo.21431874).

## Author contributions

X.S. and S.-Y.C. conceived the project. C.S. and S.R. performed data collection. M.N.W., C.S., and X.S. performed computational analyses, H.T. and M.N.W. performed experiments, Manuscript writing – Original Draft, H.T. and M.N.W.; Review & Editing, X.S., S.-Y.C., D.T., B.H., C.S., and S.R. Funding Acquisition, X.S., D.T., S.-Y.C., and M.N.W. All authors read and approved the final manuscript.

## Acknowledgements

We thank Northwestern IT Research Computing Services for providing the Quest High-Performance Computing Cluster platform and NUSeq Core for providing sequencing service. We thank Dr. Frank Furnari for providing iPSC12 cell line. We thank the Robert H. Lurie Comprehensive Cancer Center of Northwestern University for supporting Sasha Roth through the Summer Student Research Scholarship. The graphic abstract, Fig. 4A, and Fig. 5A were created with BioRender.com.

## Declaration of generative AI and AI-assisted technologies

During the preparation of this manuscript, the authors used ChatGPT to improve the readability and language of the manuscript. After using this tool or service, the authors reviewed and edited the content as needed and take full responsibility for the content of the publication.

