## Supplemental Figures for "iPSC-Derived Microglia-like Cells Exhibit Protocol-Dependent Transcriptomic Features and Robust Phagocytosis of Glioma Cells"

**Supplemental Fig. S1-2**

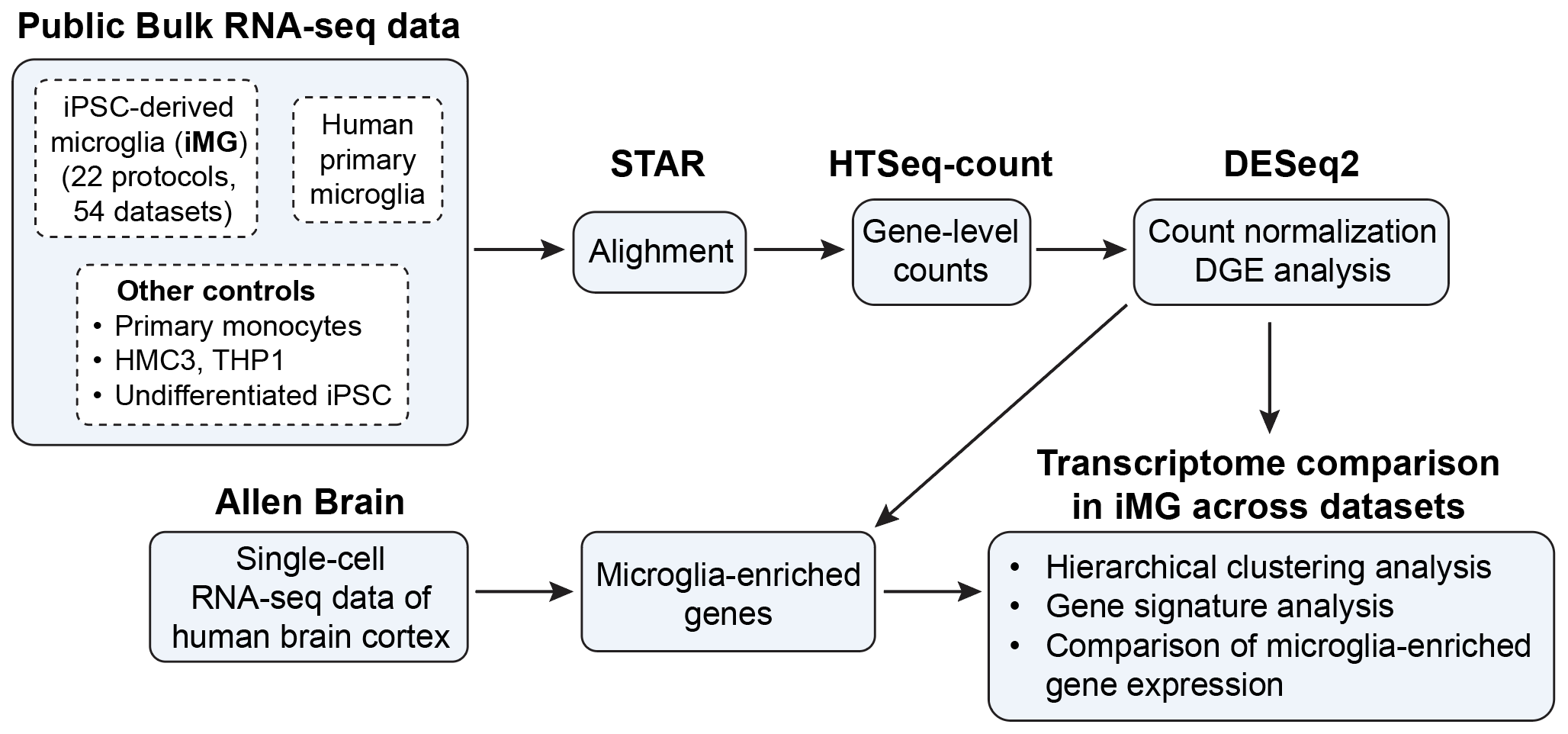

**Figure S1. Workflow of the computational analysis pipeline.**

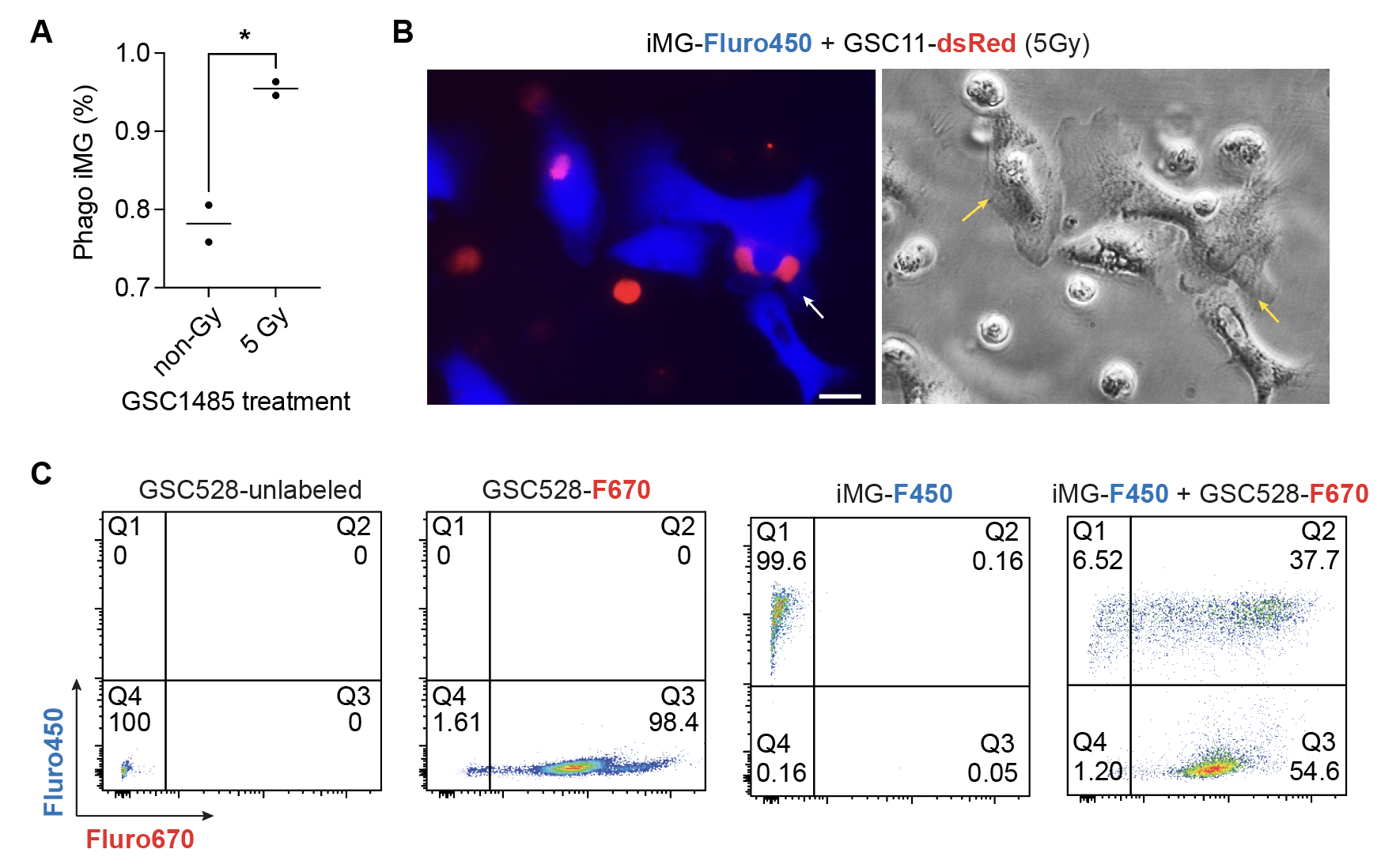

**Figure S2. EB protocol-derived iMG efficiently phagocytose glioma cells**

(**A**) Quantification of phagocytic iMG following 24-hr co-culture with GSC1485 cells under basal (non-Gy) or irradiated (5 Gy) conditions. Each dot represents an independent experiment. *, p < 0.05 (unpaired Student's t-test).

(**B**) Representative fluorescence (left) and bright-field (right) images of Fluor450-labeled iMG (blue) co-cultured with dsRed-expressing GSC11 cells (red) following 5 Gy irradiation. Arrows indicate iMG containing engulfed glioma material. Scale bar, 20 μm.

(**C**) Representative flow cytometry plots demonstrating phagocytosis of Fluor670-labeled GSC528 cells by Fluor450-labeled iMG. Phagocytic iMG were identified as Fluor450⁺/Fluor670⁺ double-positive cells (Q2) following 24-hr co-culture. GSC528 cells were cultured under basal conditions without irradiation.
